# Interplay between cell height variations and planar pulsations in epithelial monolayers

**DOI:** 10.1101/2022.05.17.492239

**Authors:** Raghavan Thiagarajan, Mandar M. Inamdar, Daniel Riveline

## Abstract

Biological tissues change their shapes through collective interactions of cells. This coordination sets length and time scales for dynamics where precision is essential, in particular during morphogenetic events. However, how these scales emerge remains unclear. Here we address this question using the pulsatile domains observed in confluent epithelial MDCK monolayers where cells exhibit synchronous contraction and extension cycles of ≈5 hours duration and ≈200 μm length scale. We report that the monolayer thickness changes gradually in space and time by more than two folds in order to counterbalance the contraction and extension of the incompressible cytoplasm. We recapitulate these pulsatile dynamics using a continuum model and show that incorporation of cell stiffness dependent height variations is critical both for generating temporal pulsations and establishing the domain size. We propose that this feedback between height and mechanics could be important in coordinating the length scales of tissue dynamics.

## I. INTRODUCTION

Cells as individual units display a variety of morphological changes during sensing [1], migration [2] and division [3] phenomena. When several individual cells are placed in close proximity to each other, each cell exhibits a complex behavior that is often different from isolated conditions [4]. This complexity further increases when hundreds to thousands of cells are organized in a continuous tissue. The collective dynamics exhibited by tissue is distinct from specific characteristics of individual cells [5]. Understanding the evolution of characteristics while transitioning from one-to-many cells is a challenge. In order to characterize these emerging properties, a combination of experimental and theoretical approaches is important to show generic principles on simple systems. In this context, spontaneous oscillations are attractive phenomena because of their clear spatio-temporal characteristics [6].

Oscillations are critical for dynamic rearrangements during the development of several organisms [7–10]. Length scales of oscillations span from molecular assemblies of proteins to group of cells in tissues [11–15]. Similarly, the periods of oscillations scale from seconds to hours [16, 17]. Several studies using experiments and theory have explored the biological pathways and the underlying molecular actors [18–20]. Although different works propose various mechanisms that set the length and time-scales associated with these pulsations, a thorough understanding of their origins is still missing.

Spontaneous planar pulsations of cells are shown to occur in Madin-Darby Canine Kidney (MDCK) monolayers [21–26]. We had previously showed [17] that pulsations appear randomly on regular culture without any coating. We also reported that these pulsations can be controlled by plating them on micro-patterns of controlled dimensions by matching pulsations sizes with grid size. Using this setup, now we show that MDCK cells also undergo height variations that inversely correlate with changes in area. We quantify the nature of these pulsations by using the power spectrum of the velocity divergence field and observe the appearance of characteristic length and time scales of ≈ 5 hours and ≈ 200 μm.

Several theoretical models, both discrete and continuum, have been developed to get insights into different aspects of tissue pulsations and wave propagation [27–29]. The discrete descriptions include active vertex [17, 26] and cellular phase field models [24], whereas the active continuum frameworks use underlying viscous or elastic rheology of the tissue to model the phenomena [23, 28]. The non-equilibrium activity is implemented in these models using cell polarity, motility, and active stress [30]. These active components are generally coupled to tissue kinematics and sometimes also with an additional chemical field such as myosin density [29]. In this paper, we developed a simplest version of a 2–D continuum model in which the tissue deformation kinematics depend on cell area and an active stress term. In such planar models, the tissue could be modeled to either be compressible or incompressible in terms of area or volume. In the current work, we model the tissue to be area compressible in 2D but incompressible in 3D along our observations of cell deformation. On the basis of this assumption of 3D volume conservation with our estimate, we also include a component to the planar tissue stress that we model to arise from spatial gradients in the thickness of the epithelial monolayer. We find that this contribution to the stress provides a part of the restoring force for pulsations and could play a critical role in setting the length scale for the pulsations.

The paper is organised as follows. In Section 2, Materials and Methods, we detail the experimental and analysis techniques used in the paper. In Section 3, Results, we present the main experimental findings of this work regarding planar pulsations and height variations in the MDCK monolayer. Here, we also develop a continuum model for tissue pulsations, and compare its findings with the experimental outcomes. Finally, in Section 4, Discussion, we provide an overview of our findings, more specifically the potential role of height variations in monolayer pulsations. We also briefly compare our approach with other related works on monolayer pulsations. Detailed derivations of the equations associated with the continuum model are provided in the Appendix.

## II. MATERIALS AND METHODS

### Cell culture

MDCK-E-Cadherin-GFP cells (Nelson lab., Stanford) were cultured in low glucose Dulbecco’s Modified Eagle’s Medium (DMEM) (31885-049, Invitrogen) supplemented with 10% FBS (10309433, HyClone) and 1% antibiotics (Penicillin-Streptomycin; 11548876, Invitrogen) at 5% CO2 in 37°C. The culture was maintained by replating cells before they reached confluence. For imaging, 1 million cells were added to 175 mm^2^ area of coverslips (CS) mounted on imaging chambers, and allowed to settle. After 1 h, the non-attached cells were washed and the media was changed to L-15 (11540556; Invitrogen) supplemented with 1% FBS. At this stage, the mean cell density was about 1 cell per 1000 μm^2^. In the next 12-16 hours following seeding, cell proliferation leads to the formation of confluent monolayer without leaving any empty space. The pulsation phase appears as soon as the monolayer is confluent and lasts about 10 hours. Later on as a consequence of cell proliferation and when the cell density is high enough to prevent large-scale movements (corresponding to a mean cell density of about 6.5 cells per 1000 μm^2^), the monolayer enters a jammed state consistent with [28, 31].

### Micro-patterning

The grid with the required dimension was fabricated as SU-8 photoresist (MicroChem) molds on silicon wafer using the standard photolithography procedure. Then stamps of PolyDimethyl Siloxane (PDMS) (Sylgard 184; Dow Corning) made with a pre-polymer to crosslinker ratio of 1: 9 (V/V) were obtained by replica molding from the silicon wafer. The micro-patterning was performed using standard soft lithography procedure as described in [17]. Briefly, coverslips were treated with “Piranha” (3:7 parts of H_2_O_2_ (516813; Sigma) and 7% H_2_SO_4_ (258105; Sigma)) for 10 min, followed by careful washing and sonication in double distilled water for 5 min. The coverslips were dried using N_2_ blower before functionalizing with (3-mercaptopropyl)trimethoxysilane (S10475, Fluorochem), and stored in a dry and clean glass petri dish at 65°C for 2 h. Meanwhile, a drop of 10 μg/ml of TRITC labelled Fibronectin (FNR01; Cytoskeleton) was added to the plasma treated PDMS stamps to allow the FN to settle on the stamp. After 1 h, the remaining liquid was removed and the stamp was carefully dried using N2 blower. The stamp was then brought in contact with the activated coverslip and left untouched for 30 minutes in order to transfer the FN to the coverslip. Finally, the coverslip was incubated with 100 μg/ml solution of poly-L-Lysine-grafted-PolyEthylene Glycol (pLL-g-PEG) (SuSoS) in 10 mM HEPES. After 20 minutes, the coverslip was rinsed three times with Phosphate Buffered Saline (PBS) (11530486; Invitrogen) and stored immersed in PBS in 4°C for up to 5 h before seeding the cells.

### Image acquisition

The fluorescent images for the height measurements were acquired using a Leica SP8 X line scanning confocal system equipped with Z galvanometric stage, controlled by LAS X interface and equipped with Oko lab stage top incubator. Using a 63x (1.4 NA, oil) or 40x (1.3 NA, oil) objective, a tile scan covering the size of a FN grid was performed with z steps of 0.5 μm. To get a larger field of view for computing the power spectrum, the phase contrast acquisitions were obtained using one of the microscopes with the corresponding objectives: inverted Olympus CKX41 (4x, 0.13 NA Phase, Olympus); SANYO MCOK-5M incubator microscope (10x, 0.25 NA Phase, Olympus). These microscopes are equipped with a CCD camera (Hamamatsu C4742-95 / C8484-03G02; Sentech XGA) and operated by Hamamatsu Wasabi / Micromanager and MTR-4000 softwares respectively. The interval between acquisitions were either 5 min, 10 min or 20 min and all the imaging was performed at 37°C for 48 h. To prevent evaporation of media during imaging, either mineral oil (M8410, Sigma Aldrich) was added or the imaging chamber was covered with a glass Petri dish.

### Image processing

The images were analyzed using FIJI application [32] and the power spectra were computed using Matlab software. The schemes were drawn using Inkscape and the plot showing the difference in height were obtained using Graphpad Prism 7. The height variations in the monolayer were obtained using a custom written FIJI plugin. The plugin allows visualization of changes in height as variations in intensities by encoding the height as intensity value to the corresponding pixel. In order to get the power spectrum of the pulsations, the velocity field was obtained by Particle Image Velocimetry (PIV) using the Matlab based PIVlab application [33]. All experiments were repeated a minimum of three times and the typical results are reported.

## III. RESULTS

In this section, we present the main findings of the current work on (i) localisation of planar pulsations on patterned substrates, (ii) the correlation of height variation in the monolayer with planar contraction/expansion, (iii) quantification of the length and time scales associated with the monolayer pulsations, and (iv) a continuum model that recapitulates experimental results.

### A. Planar pulsations in MDCK monolayers are localised on a substrate patterned with fibronectin and PEG

We prepared MDCK monolayers with controlled density on coverslips with patterns of fibronectin (FN) (see Materials and Methods and fig. 1). These adhesive patterns were grids of 120 μm thickness surrounding a square gap of 150 μm, passivated by pLL-g-PEG (PEG) that reduces adhesion of cells to the coverslips (fig. 1). This setup generates a region of low friction connected to zones of high friction. As we previously reported [17], the FN grid led to spontaneous pulsations of MDCK cells spanning 10-15 cells in diameter. To gain insight into cell shape changes in 3D, we imaged the pulsating domain over 48 h using confocal microscopy (fig. 2a). This experiment allowed us to gain insight into the potential correlation between the change in cell height and pulsations.

**FIG. 1.**
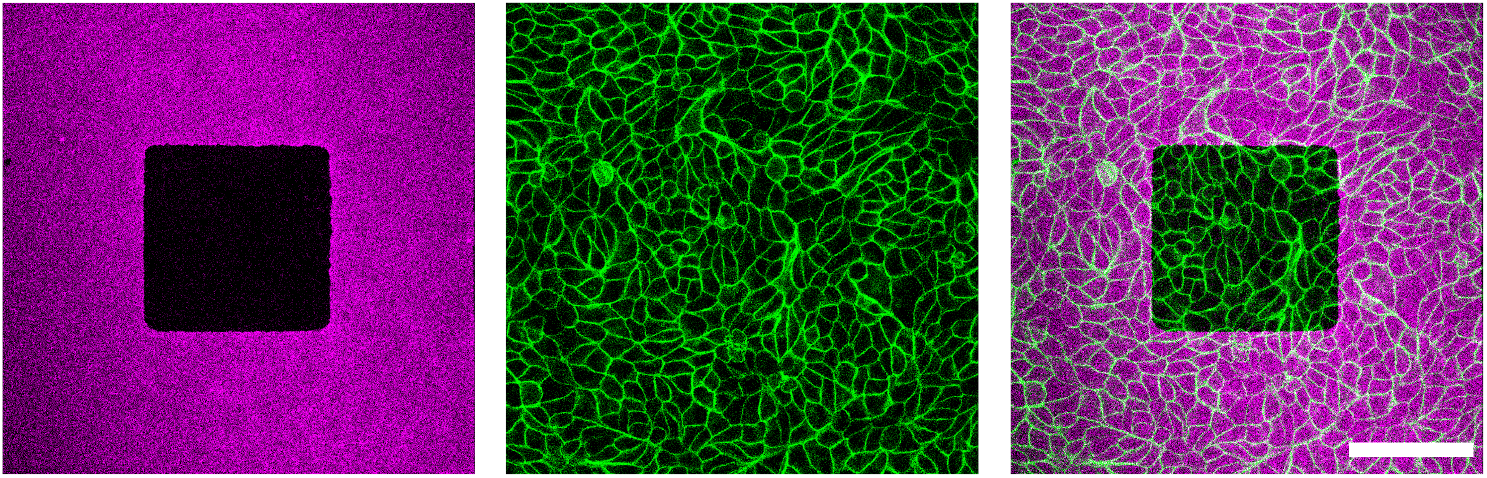
Visualizing a domain of MDCK monolayer. The first image shows the FN grid of width 120 μm and gap of 150 μm. The second image shows the MDCK-E-Cadherin-GFP cells in green and the third image shows the overlay of cells on FN grid. Scale bar, 100 μm.

**FIG. 2.**
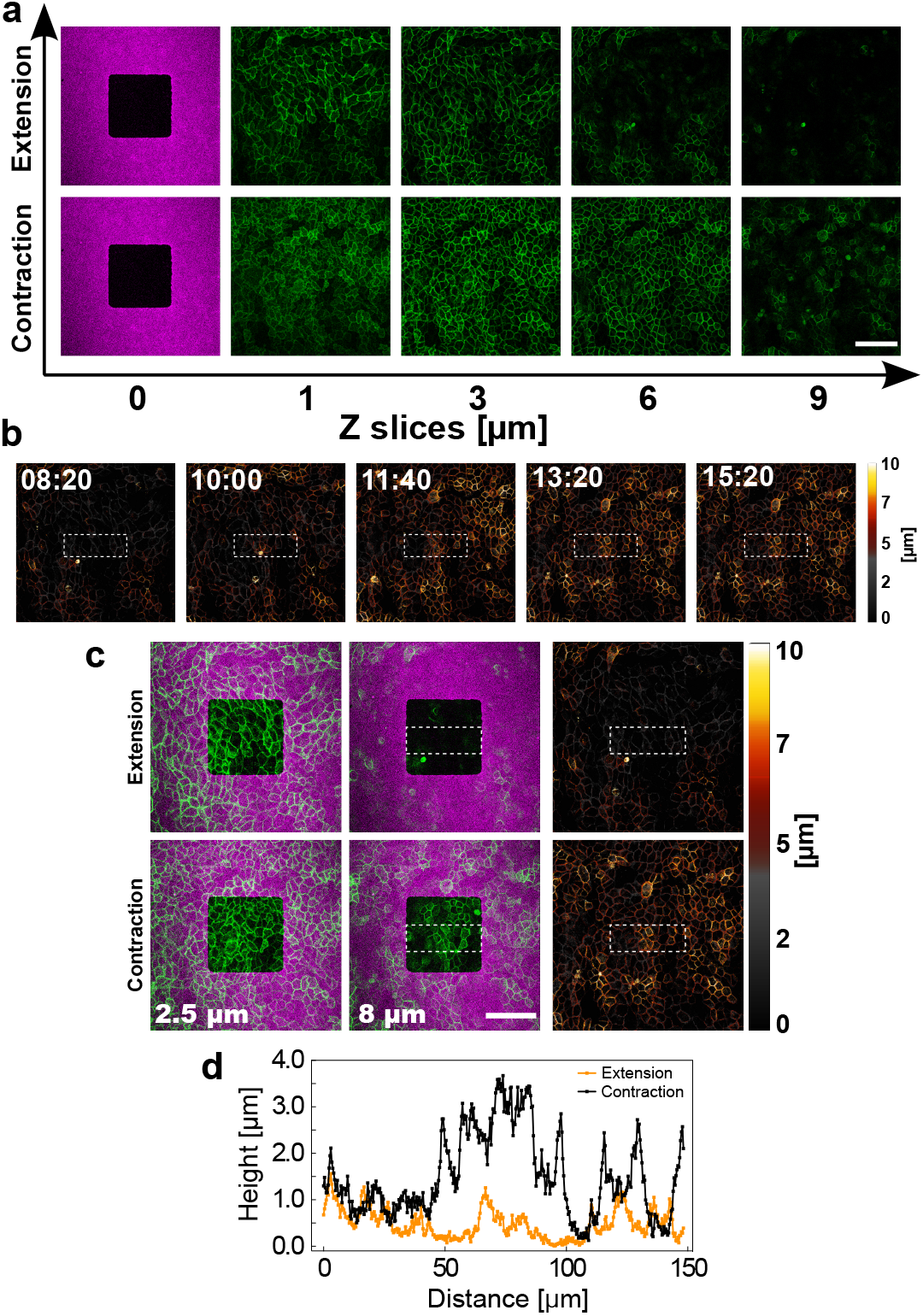
MDCK cells imaged at different heights (confocal z-planes). (a) The two rows of images correspond to the extension and contraction phases. Images are arranged from left-to-right indicating the appearance of cells at increasing heights. The FN grid on the coverslip surface is taken as the reference and therefore is denoted as 0 μm. Majority of the cells are visible at higher Z planes only in the contraction phase. (b) - (d) Analysis of height fluctuations. (b) Snapshots of images showing the height of cells encoded as intensity, corresponding to the domain shown in (a). The first image (08:20) shows the extension phase and the last image (15:20) shows the contraction phase. Time is in hh:mm. (c) The first and second rows represent the extension and contraction phases of the domain shown in (a). The first two columns correspond to different heights (2.5 μm and 8 μm) to show the differences in the appearance of cells during the extension and contraction phases. The last column shows the images of cells where the height is encoded as intensity values with higher intensities representing higher z-planes. These images are the same as shown in (b) at 08:20 and 15:20 respectively. (d) Plot showing the mean height of cells corresponding to the dotted rectangle in the images of last column in (c). The mean intensity values are obtained by taking the plot profile of the rectangle, where the intensity is represented as one (averaged) value for every column along the length of the rectangle. The rectangle length is denoted as d7istance in the x axis and the mean height of cells are denoted by the y axis. In (a) and (c) the MDCK-E-Cadherin cells are shown in green and the FN grid is shown in magenta. Scale bar, 100 μm. Color bar in (b) and (c) indicates the corresponding height values. Also see Movie 1.

### B. Pulsations in MDCK monolayers correlate with variations in cell height

During the pulsations, we found that the height of cells in the monolayer underwent fluctuations. This was confirmed by checking the appearance of apical side of cells at different axial planes over time (fig. 2a). During the contraction phase, cells were visible already at z-plane 8 μm, while in the extended phase cells were visible only at the z-plane 2.5 μm (fig. 2c). The maximum height reached by cells in the monolayer was about 10 μm (fig. 2a). In addition, to quantify the height profiles, we encoded the cell height as intensity values (fig. 2b and 2c and (movie 1)). The results confirmed our visual observation and the mean height profile of cells measured over a region of interest across time, showed variations in mean cell height (fig. 2b and 2c). Specifically, the height profiles between the contraction and extended phases were more than two-fold different (fig. 2d). This confirmed that pulsations are correlated to cell flattening with reduced height during the extension phase followed by cell squeezing with increase in height during the compression phase. The Cell volume was estimated to be roughly constant, at around 400 μm^3^.

### C. Planar pulsations in MDCK monolayers have characteristic time and length scales

As described earlier, the pulsations observed in the epithelial monolayer corresponded to the periodic formation of contracting and expanding domains within the monolayer. To quantify the pulsations, we first used particle image velocimetry (PIV) to obtain velocity field **v**(*x*, *y*, *t*) of the cells at grid points *x_i_, y_j_* at time-frame *t_k_*. We then numerically obtained the planar divergence *d*(*x, y, t*) = ∇ · **v** since it is the relevant measure of contraction/expansion in the monolayer due to the velocity field. Representative divergence fields in space at a particular time and at a given space point as a function of time are shown in figs. 3a and 3b, respectively. In order to extract the spatiotemporal characteristics of divergence, we numerically obtained its power-spectrum *S_d_*(*q_x_,q_y_, ω*) through discrete Fourier transform where *q_x_* and *q_y_* are the wave-numbers, respectively, along *x* and *y*, and *ω* is the frequency component of the power spectrum. To quantify the temporal behavior of *d*, we then calculated *S_d_*(*ω*) by summing *S_d_*(*q_x_,q_y_, ω*) over all *q_x_* and *q_y_*. The spatial characteristics of *d* are quantified via *S_d_(q)* by summing *S_d_*(*q_x_, q_y_, ω*) over all *ω* and also over *q_x_* and *q_y_* such that 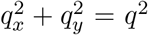. Note that for convenience, we keep the same notation *S_d_*, irrespective of the associated argument. The power-spectra *S_d_*(*ω*) and *S_d_*(*q*) are plotted in figs. 3c and 3e, respectively. The blue line and the surrounding gray region correspond, respectively, to the average and the standard deviation of *Sd*(*ω*) and *Sd*(*q*) over three separate experimental repeats. It can be seen that *S_d_*(*ω*) has a peak at *ω*/2*π* ≈ 0.2 h^-1^ that corresponds to a pulsation period of ≈5 h. The peak corresponding to *S_d_*(*q*) is not as sharp and also shows some variation with respect to the experimental repeats. However, *S_d_*(*q*) peaks at *q*/2*π* ≈ 0.005 *μ*m^-1^ which corresponds to a periodic pattern in divergence with a length-scale of ≈200 *μ*m.

**FIG. 3.**
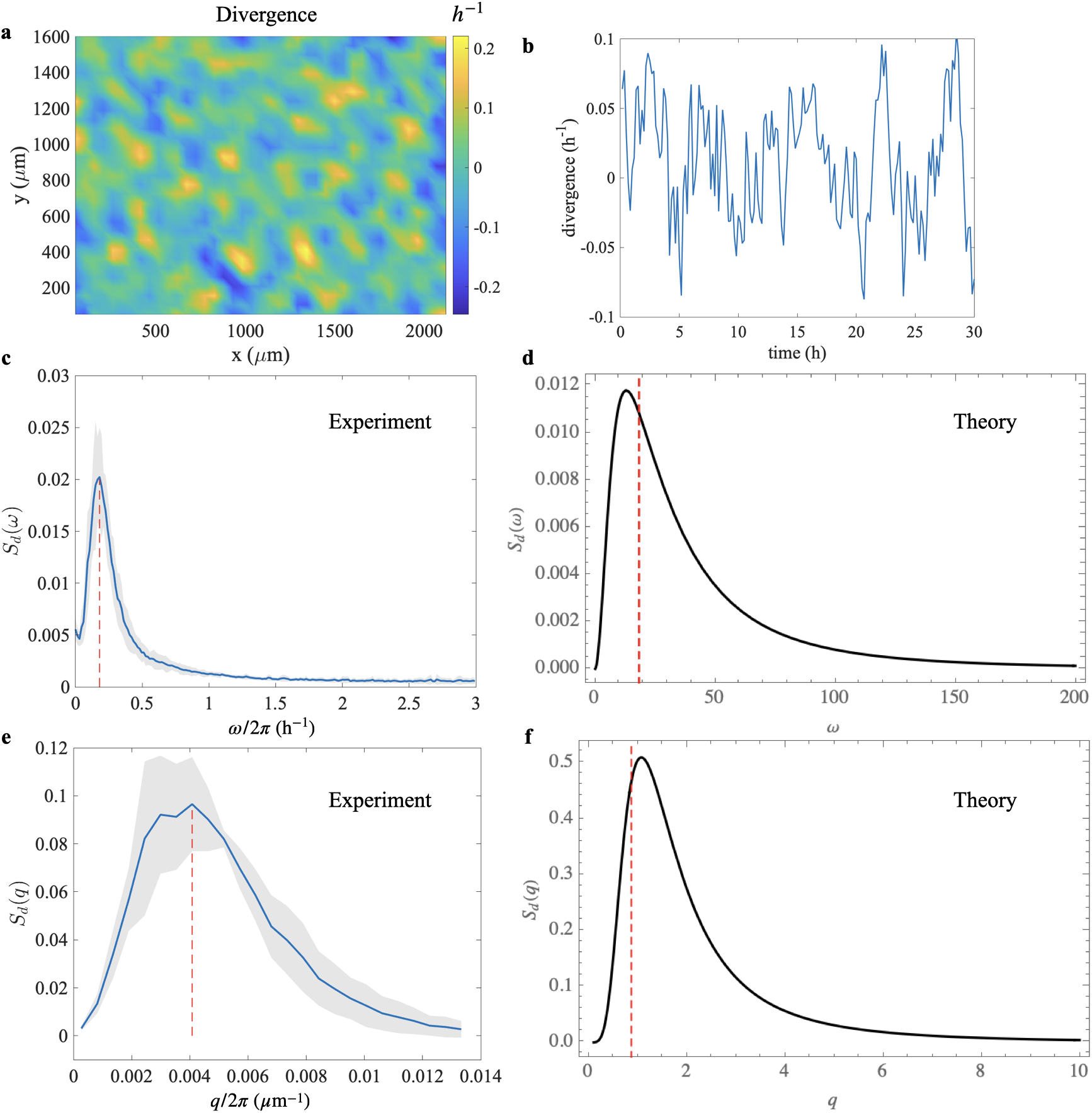
Representative divergence field obtained from velocity field of one experimental sample in (a) space at a given instance in time and (b) with respect to time at a fixed space point. The regular pattern in space and time is visible and systematically quantified using Power spectrum density *S_d_* as a function of wave number *ω* (c) and frequency *q* (e) obtained from the combined experimental data. The blue lines and the gray shaded region correspond to the mean and standard deviation over three separate experimental repeats. The red lines correspond to the location of the peak values of the spectrum. The equivalent theoretical calculation for *S_d_*(*ω*) (d) and *S_d_*(*q*) (f), in non-dimensionalized units, are also shown. The red lines in the theoretical plots correspond to the rough estimate of the peak location *ω_m_* from Eq. A29 and *q_m_* from Eq. A26. The non-dimensional parameters (eq. 7) used for the theoretical curves are *K* = 0.1, *K_h_* = 40, *τ* = 0.1, *μ* = 2, *τ_c_* = 0.02.

Thus, we observe from the experiments that MDCK monolayers undergo planar pulsations with characteristic time and length scales. We also found that the planar pulsations were accompanied with modifications in the monolayer thickness that was both space and time dependent. Moreover, it was also shown in our earlier work that gradients in myosin concentration were involved in monolayer pulsations [17]. Hence, in order to get a better understanding of how planar deformations, thickness modifications and contractile mechanisms together contribute to the observed spatiotemporal pulsatory patterns in epithelial monolayers, we developed a continuum model based on a previous work [34].

### D. Continuum model for spontaneous pulsations in MDCK monolayer

The experimentally observed pulsations in epithelial monolayers could be quantified in terms of the velocity divergence of the tissue flow that captures the dynamics of area expansion and contraction of tissue domains. It was also observed earlier [17] that myosin contractility was associated with these collective deformation modes of the monolayer. Hence, we now develop a simple continuum model with cell area *a*, contractile matter *ζ*, and velocity field *v* as three variables that are functions of *x, y*, and *t*. The mass conservation equation can be written as

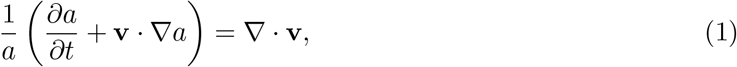

by neglecting the contributions from cell division and death. The stress *σ_ij_* within the tissue should satisfy the momentum balance equation that is given by

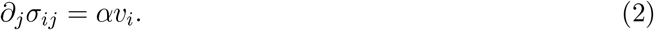

Here, we assume that the imbalance in the internal tissue stress is countered by the passive fluid friction between the tissue and the substrate. Cell-substrate interaction could also produce active substrate traction, possibly arising from cell motility. However, for simplicity, in our model we do not include this potential contribution. In the above equation as well as the ones below, we use indicial notation for vectors and tensors with the indices *i, j* representing *x, y* coordinates. Since we are concerned only with the contractile and expansive movements of the tissue, the stress is modeled as an isotropic quantity

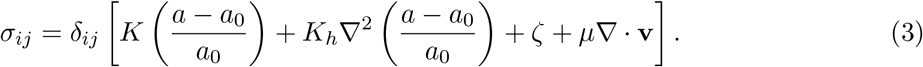

Here, *a_0_* is the homeostatic area of the cells, *K* is the monolayer area modulus, *K_h_* is the gradient elasticity modulus that energetically penalizes the height variations in the monolayer (also see Appendix C), and *ζ* is the myosin dependent contractile active stress in the tissue. Finally, we discuss the dynamics of the contractile material *c*(*x*, *y*, *t*). For simplicity, we model the active stress *ζ* = *Bc*, where *B* is a proportionality constant. Consequently, to prevent the introduction of a new field variable *c*(*x,y,t*), we simply replace *c* with *ζ* without losing generality. Hence the dynamics of the active stress *ζ* in the tissue is given by

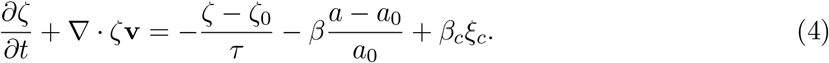

Note that the use of *ζ* instead of *c* would involve a simple linear scaling of the parameters *β* and *β_c_* with the factor *B*. Here, *τ* is the time-scale for *ζ* to reach its homeostatic value *ζ*_0_, *β*>0 is the coupling term that modulates *ζ* depending on deviation of cell area from *a*_0_, and *ξ_c_* is the chemical noise with strength *β_c_*.

We then combine the above equations and linearize them for small perturbations *δ_a_* = *a* – *a*_0_ and *δζ* = *ζ* — *ζ*_0_. We then replace 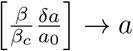 and 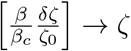, and non-dimensionalise length and time with 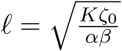 and 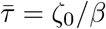, respectively. Finally, we get the following non-dimensionalised equations (see Appendix A for details)

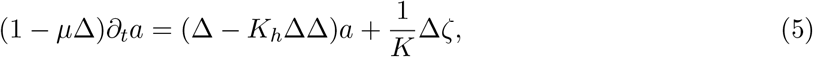

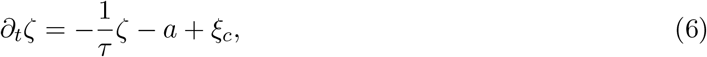

Here, the non-dimensionalisation resulted from the following substitutions of parameters in terms of the original parameters.

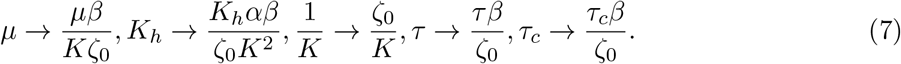

Without the noise term *ξ_c_*, the above set of equations are satisfied for *a* = 0 and *ζ* = 0, and result in *v_i_* = 0. However, for non-zero noise, the system is perturbed and the feedback between *a* and *ζ* results in persistent fluctuations having dominant length and time scales that, as we show below, depend on the tissue parameters. Thus in our model, the noise plays an essential role in sustaining the pulsatory movements. By modeling *ξ_c_* to be correlated in time and uncorrelated in space (see Appendix A), and doing Fourier analysis of these set of equations, the power-spectrum *S_d_*(*q_x_*, *q_y_*, *ω*) of velocity divergence can be obtained exactly. Further, *S_d_*(*q,ω*) = 2*πqS_d_*(*q_x_,q_y_, ω*), where 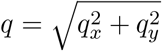, from which

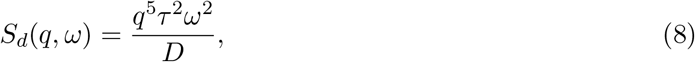

and the denominator

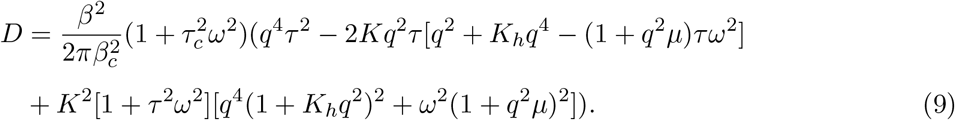

The noise term *ξ_c_* can have its origin, for example, in the modulation in myosin levels at cell-cell contacts that are known to cause junctional fluctuations [35, 36].

The theoretical estimates of *S_d_*(*ω*) and *S_d_*(*q*) are presented in figs. 3d and 3f, respectively and show qualitative similarity with their experimental counterparts that include the presence of peak in *S_d_*(*ω*) and *S_d_*(*q*), thus providing characteristic time and length scales for the pulsatory cellular movements. Moreover, as is experimentally observed, both *S_d_*(*q*) and *S_d_*(*ω*) decay to zero for larger values of *q* and *ω*, respectively. However, unlike for the experimental data in which *S_d_*(*ω* = 0) ≠ 0 (fig. 3c), the theoretically estimated value is zero (fig. 3d). By performing simple asymptotic analysis on *S_d_*, the dominant length (*l*_0_), in terms of the original parameters, can be calculated as

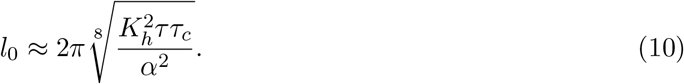

giving the approximate location of the peak for *S_d_*(*q*) in fig. 3f (see Eqs. 7 and A26). We obtain *l*_0_ of about 100 *μ*m, by taking typical values reported or estimated for the four parameters (see Appendix D). It can be seen that for a pulsating length scale to emerge in the monolayer, *K_h_*, the gradient modulus that is associated with spatial height variation is necessary as per our model. Similarly, the other important quantities for the emergence of pulsation length scale are the turnover timescale *τ* for myosin and the time-scale *τ*_c_ for the chemical noise. It can be seen that if any of these quantities tend to zero, then the peak of *S_d_*(*q*) is pushed towards higher values of *q* and eventually disappears. Moreover, when *τ_c_* → 0, i.e., the chemical noise is uncorrelated or white, the decay in *S_d_*(*q*) and *S_d_*(*ω*) is much slower than is experimentally observed. Thus the qualitative behavior of the spatiotemporal patterns exhibited by the tissue in our model depends on a combination of tissue mechanical properties and characteristics of the chemical noise. We point out that the amplitude of the tissue response would be proportional to the noise amplitude *β_c_*. However, since we plot only the normalized values of power-spectra, the noise amplitude does not explicitly show up in figs. 3d and 3f. The details of this calculation and other estimates for the power-spectrum including the experimental comparison are provided in Appendix A. Based on these estimates and additional numerical exploration, we find that the qualitative behavior of *S_d_*(*q*) and *S_d_*(*ω*) (figs. 3d and 3f) is similar to their experimental counterparts (figs. 3c and 3e) for a wide range of parameters. We chose the typical parameters that showed this trend (fig. 3 caption). We note that although the experimental data (figs. 3c and 3e) are obtained for the monolayer patterned substrate as shown in fig. 1, the theoretical model does not account for the variations in substrate frictions that are expected to be inherent in the experiments. Hence, we also calculated *S_d_*(*ω*) and *S_d_*(*q*) from our previous data on non-patterned substrate [17]. We find that the differences between *S_d_*(*ω*) and *S_d_*(*q*), especially with respect to the location of maxima in *q* and *ω*, are not significant (see Appendix B).

## IV. DISCUSSION

We show that coordinated cell height variations with the planar contraction and extension cycles is potentially important in determining the size of pulsatile domains in monolayer tissues. Shape fluctuations in biological systems range from cell membrane [37] and organelles [38] to single [13] and collection of cells in embryos [15]. During developmental phenomena in which motions of cell collections can be synchronized, such movements underlie crucial morphodynamic events that exhibit outstanding spatial precision [39]. In this context, our experimental findings suggest that variations in cell height are associated with changes in cell area in a nearly incompressible viscoelastic cytoplasm, and are consistent with one of our model assumptions. The coupling between the contractile machinery and the cell area, highlighted by *β* in Eq. 4, indicates that due to the chemical noise *β_c_* (Eq. 4), the concentration of the contractile material *ζ* can change on a cell by cell basis leading to varying levels of tension and therefore inhomogeneous height distributions across the pulsatile domains. The variations in height profiles of cells indicate that the height of cell-cell junctions is shortened and extended, respectively, during the extension and contraction phases. Therefore, the contractile stress of the cells combined with cell-cell junction height diminishes away from the pulsation centre. In this specific context, the link between junction height and the contractile molecules is an interesting question to be explored, since the accumulation of myosin could be due to intercellular flows [40] or due to transcriptional activity [41]. Finally, it would be interesting to analyse the height variations with varying cell density at plating, since cell density was reported to have direct effects on cell velocities [28] and cell extrusion [31].

In our model, the relaxation of the contractile stress, dictated by the turnover time *τ* (Eq. 4), correlation time of chemical noise *τ_c_*, height gradient stiffness modulus *K_h_* (Eq. 3), and substrate friction α are crucial in dictating the inherent size *l*_0_ of the pulsating domain of the monolayer (Eq. 9). Interestingly, although *l*_0_ does not depend on the contractile activity *ζ*_0_ in the tissue, the frequency of pulsation depends on a combination of the active stress and mechanical properties of the tissue (Eq. 10). All these quantities can be experimentally measured for the monolayer under the given experimental conditions. For example, the turnover time can be measured by Fluorescence Recovery After Photobleaching (FRAP) of myosin in cells at different positions on the pulsating domain. The model can be further extended to explicitly include the role of signalling pathways that regulate contractility and deformation dynamics in tissues [36]. In addition, although we ignored the role of cell division and apoptosis on tissue area fluctuations, myosin accumulation that is involved in these process could be important in regulating the dynamics of pulsatile domains in the monolayer [42]. Indeed, not accounting for some of these important quantities and coupling could potentially explain the discrepancy between the experimental values and theoretical predictions (Appendices A and B, figs. 3–5).

Interestingly, other studies report the origin of pulsations to osmotic pressure from water flux where MDCK cells do not change significantly in height during pulsations [21]. Similarly, in single cell zebrafish embryos, bleb formation which is primarily due to contractility is also regulated by water flow through aquaporins [43]. *In vivo,* during dorsal closure by amnioserosa cells in *Drosophila,* the force required for contraction has been shown to arise from both contractile actomyosin cable and cell volume change regulated by potassium channels [44]. Along this line, our study can be further extended to volume measurements (from apical-basal area and cell height) to determine the respective contributions of contractility and osmotic pressure in pulsations in MDCK monolayers. Such precise measurements can be used to refine the hydrodynamic descriptions for tissue dynamics with different sources of stresses and material flux [45]. Together, this will lead to detailed insights on the role of mechanics in tissue dynamics and organisation at different scales.

## ACKNOWLEDGMENTS

The MDCK-GFP-E-Cadherin cells were a kind gift from Nelson lab., Stanford. We are grateful to Guillaume Salbreux for discussions and many key insights into the theoretical model. We thank the Riveline lab. for discussions and comments, and acknowledge the help of Basile Gurchenkov, Yves Lutz and Pascal Kessler from the IGBMC imaging facility. We thank the late Marcel Boeglin for writing the Fiji plugin to perform height measurements. R.T. was an IGBMC international PhD program fellow supported by LabEx INRT funds. M.M.I acknowledges funding from Science and Engineering Research Board (MTR/2020/000605). D.R. acknowledges support from CNRS (ATIP), ciFRC Strasbourg, the University of Strasbourg, Labex IGBMC. This study has been also supported by a French state fund through the Agence Nationale de la Recherche under the frame program Investissements d’avenir labeled A NR-10-IDEX-0002-02.

## Author contribution

All authors contributed equally to this paper.

## Appendix A: Derivation of equations for the continuum model and analytical estimates

Here, we systematically develop the continuum model described in the main paper, and also provide detailed derivations of the relevant equations.

The theory has 2D cell area *a*, velocity field of the cells *v_i_*, internal stress *σ_ij_*, and the contractile matter *ζ* as the field variables [34]. The relevant equations are

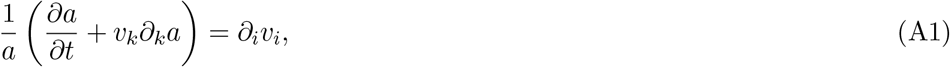

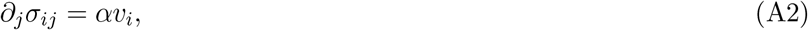

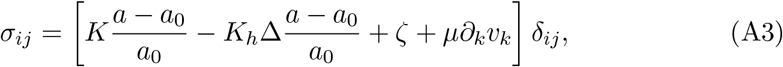

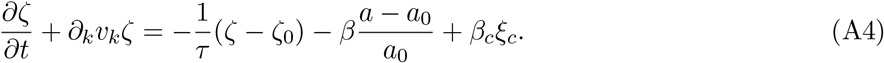

In the above equations, *α* is the friction coefficient between the monolayer and the substrate, *K* is the linear area modulus of the cells, *K_h_* is the height modulus of the tissue that is modeled to depend on the area gradient in the tissue (see Appendix C). In Eq. A4, *τ* is the time required for myosin to reach its homeostatic value *ζ*_0_, *β* > 0 is the coupling coefficient that relates cellular area with *ζ* modulation, and *ξ_c_* is time correlated noise satisfying

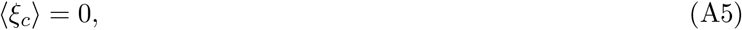

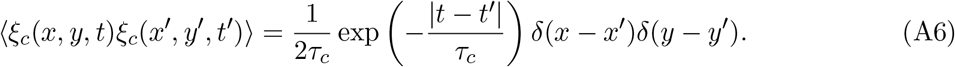

In the limit when the correlation time *τ_c_* → 0 the noise becomes uncorrelated or white noise. The parameters in the continuum equations and their units in SI are summarized in Table 1 - see also below in Appendix D.

**Table I.** SI units of variables and parameters associated with the model

| Quantity | SI unit |
| --- | --- |
| $a$ | $\text{m}^2$ |
| $\zeta$ | $\text{N/m}$ |
| $\zeta_0$ | $\text{N/m}$ |
| $v$ | $\text{m/s}$ |
| $K$ | $\text{N/m}$ |
| $K_h$ | $\text{N} \cdot \text{m}$ |
| $\mu$ | $\text{N} \cdot \text{s/m}$ |
| $\alpha$ | $\text{N} \cdot \text{s/m}^3$ |
| $\tau$ | $\text{s}$ |
| $\tau_c$ | $\text{s}$ |
| $\beta$ | $\text{N} \cdot \text{m}^{-1} \cdot \text{s}^{-1}$ |
| $\beta_c$ | $\text{N} \cdot \text{m}^{-1} \cdot \text{s}^{-1}$ |

For small perturbations, *δa* = *a* – *a*_0_ and *δζ* = *ζ* – *ζ*_0_, eliminating the velocity term, and linearizing in *δa* and *δζ* we get

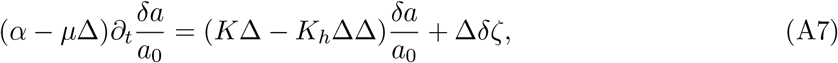

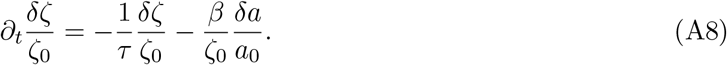

Here, we neglect the advection term of the form *ζ*_0_*∂_k_v_k_* for conceptual simplicity in the weak advection limit. Choosing, 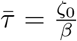 as a reference time scale and 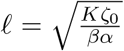 as the length-scale, we get the following non-dimensionalized dynamical equations.

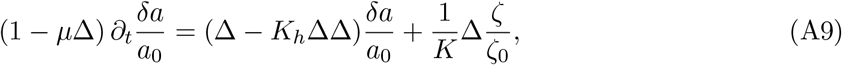

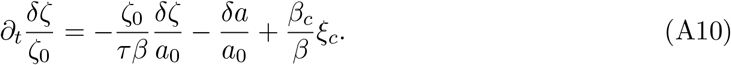

We can further re-write these equations to give

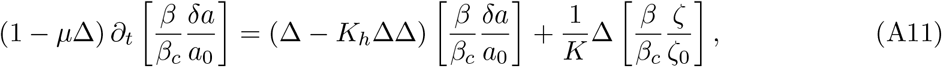

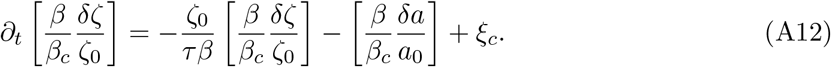

Finally, we present the governing equations as

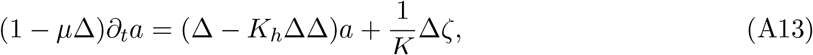

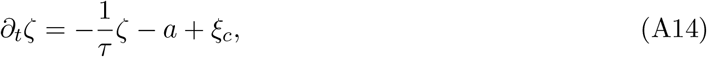

where the non-dimensional dynamical variables are written as 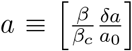 and 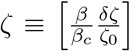. The four, free, non-dimensional parameters in these equations are 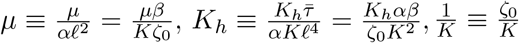, and 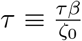, in terms of the original dimensional parameters. The correlation time of the colored noise *ξ_c_* is written as 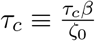.

Taking Fourier transform of the two coupled equations in space (*x* → *q_x_* and *y* → *q_y_*) and time (*t* → *ω*), we get

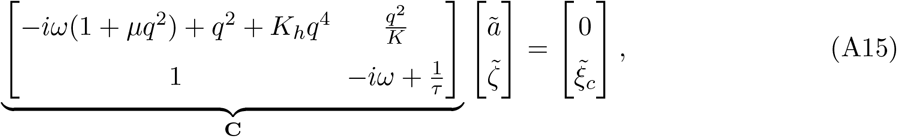

where *ã* and 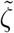 are the space-time Fourier transforms, respectively, of *a* and *ζ*, and 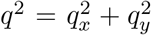.

Inverting the **C** matrix, we get

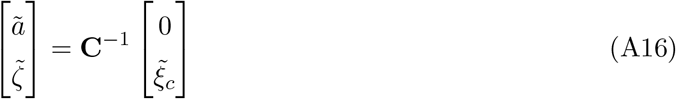

The power spectrum from ã and 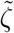 then becomes

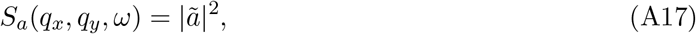

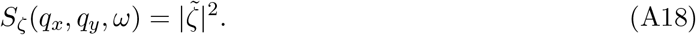

We note that since 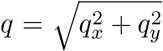, the power spectrum only in terms of *q* can now be obtained as *S_a_*(*q,ω*) = 2*πq* × *S_a_*(*q_x_, q_y_, ω*). Since the linearized mass balance equation Eq. A1 in terms of the non-dimensionalised variables and parameters is

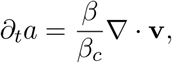

the power spectrum for velocity divergence becomes, 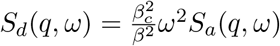. Since we have taken *ξ_c_* to be correlated noise (Eq. A4), its power-spectrum 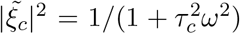. Hence, the effective power spectrum for the velocity divergence is given as

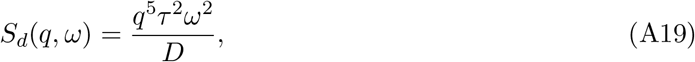

where

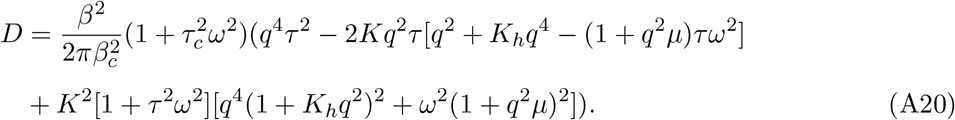

We can perform a scaling near *q* → ∞

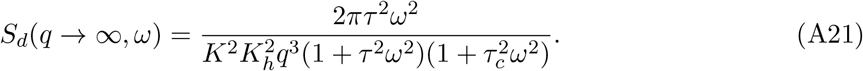

Upon integrating the *ω* terms out we get

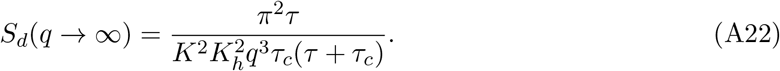

and near *q* → 0 we get

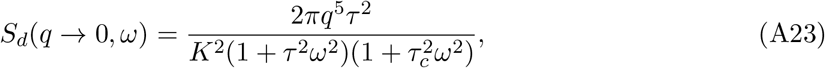

upon integrating the *ω* terms we get

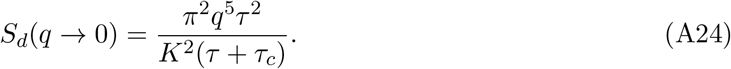

As derived above, the series expansion for *S_d_*(*q*) is increasing and decreasing in *q* for small and large values, respectively, of *q*. Hence, we could expect the location *q_m_* of the maxima of *S_d_*(*q*) to approximately be at the intersection of these two curves. Hence, we equate them

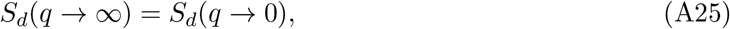

to give

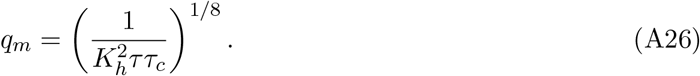

The wavelength associated with this pattern will be given by *l*_0_ = 2*π*/*q_m_*.

Following the same procedure as for *q*, we now check for the asymptotic behavior for *S_d_*(*q, ω*) in terms of *ω*.

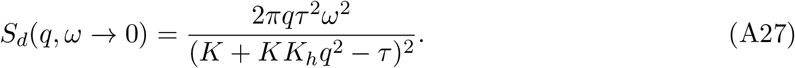

Similarly

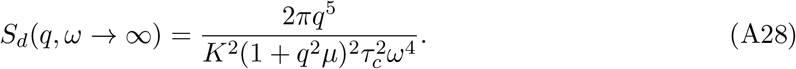

We find that the location of maxima for *S_d_*(*ω*) is obtained quite well when we equate *S*(*q,ω* → 0) and *S*(*q*, *ω* → ∞) corresponding *q* = *q_m_* as obtained in Eq. A26. From that we get that the maxima for *S_d_*(*ω*) occurs at

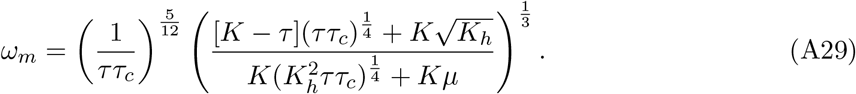

Based on Eqs. A21-A29, we can get some insights into the role the non-dimensionalised parameters *K*, *K_h_*, *μ*, *τ*, and *τ_c_* on the behavior of power spectra *S_d_*(*ω*) and *S_d_*(*q*). From Eq. A26, we find that, as per our model, the three terms *K_h_*, *τ*, *τ_c_* are important for the observation of maxima in *S_d_*(*q*) as is observed experimentally (fig. 3e). Hence, the term corresponding to gradients in area in tissue stress (Eqs. 3 and A3), the active stress homeostatic term (Eqs. 6 and A14), and the time-correlation component of the noise (Eq. A6) play key role in setting the length scale of pulsations in the tissue within our model framework. Additionally, from Eqs. A22 and A24, we see that the rate at which *S_d_*(*q*) grows and decays for small and large values, respectively, of *q* is governed by the parameters *τ*, *K*, and *K_h_*. From Eq. A28 it can be seen that although the parameter *μ* does not play an essential role in governing the qualitative nature of *S_d_*(*q*) and *S_d_*(*ω*), it modulates the decay behavior of *S_d_* (*ω*) for larger values of *ω*.

In fig. 3, we plot the normalized power spectrum in terms of the wavenumber *q* and *ω* by defining the following.

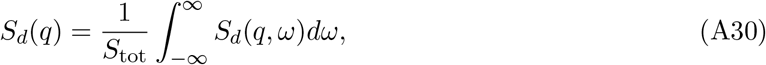

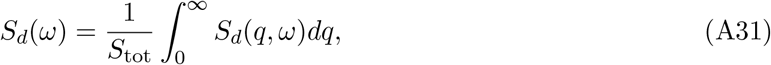

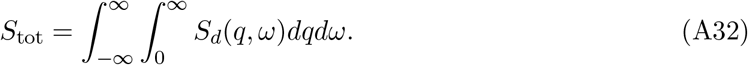

Note that for notational simplicity, we use the same symbol Sd for the different variants of the power-spectrum.

In order to quantitatively obtain the low and high *q* and *ω* behavior of *S_d_*(*q*) and *S_d_*(*ω*), respectively, we show in fig. 4 log-log plot of the data presented in figs. 3a and 3c. Since, as opposed to the theoretical calculation, *S_d_*(*ω* = 0) ≠ 0 for the experimental data, we fit a power-law of the form *ω^m^* while neglecting a few data points at the beginning (fig. 4a). The exponent *m* ≈ 1.4 and *m* ≈ −1.7 (fig. 4a) is not the same as *m* ≈ 2 (Eq. A27) and *m* ≈ −4 (Eq. A28), respectively, as is predicted by the model. Similarly, the exponent *m* ≈ 1.5 and *m* ≈ −4.8 (fig. 4b) for *S_d_*(*q*) also does not match with the theoretical estimate of *m* = 5 (Eq. A24) and *m* = −3 (Eq. A22).

**FIG. 4.**
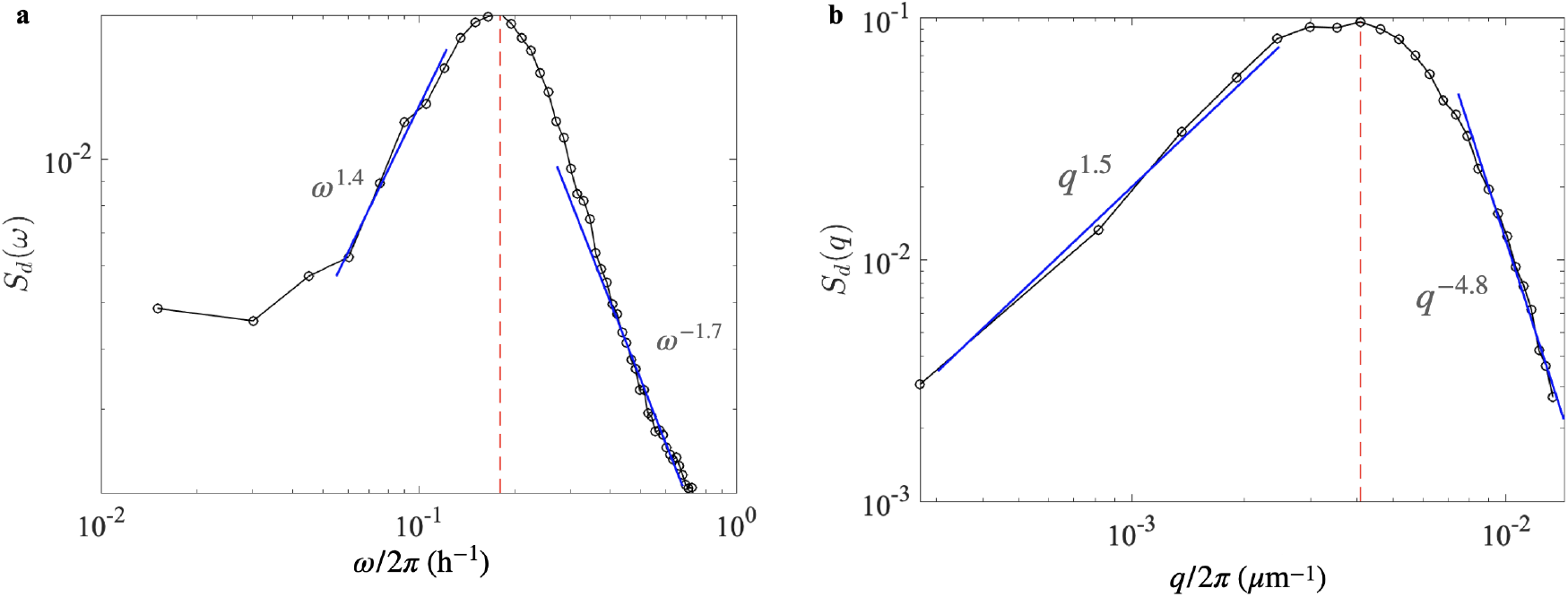
Log-log plot for (a) *S_d_*(*ω*) and (b) *S_d_*(*q*) presented earlier in Figs. 3c and 3e, respectively. The behavior of the power-spectrum at small and large values of *ω* and *q*, respectively, are shown as blue lines in (a) and (b). (a) For small values of *ω*, *S_d_*(*ω*) ~ *ω*^1-4^ and for larger values of *S_d_*(*ω*) ~ *ω*^-1-7^. The theoretical values of these exponents, 2 (Eq. A27) and –4 (Eq. A28), are not the same as those experimentally observed. (b) For small values of q, S_d_(q) ~ q^1-5^and for large values of q, S_d_(q) ~ q^-4-8^. The theoretical values of these exponents, 5 (Eq. A24) and –3 (Eq. A22) are different than their experimentally obtained counterparts.

## Appendix B: Power spectrum for cellular pulsations on non-patterned surface

In our previous work [17], we had studied pulsations on epithelial monolayers. Here, similar to the power spectrum *S_d_*(*ω*) and *S_d_*(*q*) for the divergence of pulsatory cellular movements on patterned surfaces discussed in Section 3.3 and plotted in figs. 3c and 3e, respectively, we do the equivalent for the non-patterned surfaces in figs. 5a and 5c. As shown in fig. 5, the blue curves and the gray shaded region correspond to the mean and standard deviation of the quantities over five experimental repeats. The peak for *S_d_*(*ω*) happens at *ω*/2*π* ≈ 0.15 h^-1^ corresponding to oscillation period of approximately 6.5 h. Although the peak is not as well defined for *S_d_*(*q*), its maxima occurs at around *q*/2*π* ≈ 0.004 *μ*m^-1^ corresponding to a periodic pattern of approximately 250 *μ*m. The values are not significantly different from that obtained for the patterned surface. We also check the values of the power spectra at larger and smaller values of *ω* (fig. 5b) and *q* (fig. 5d) and obtain the power-law exponents as was done in fig. 4 for patterned substrates. Although the qualitative rise and decay in *q* and *ω* for the power-spectra is similar for the patterned and non-patterned surfaces, the power-law exponents are not the same for these two conditions. However, the exponents for *S_d_*(*q*) do exhibit a better match with each other. As for the patterned surface, the exponents also do not match the values predicted by the model.

**FIG. 5.**
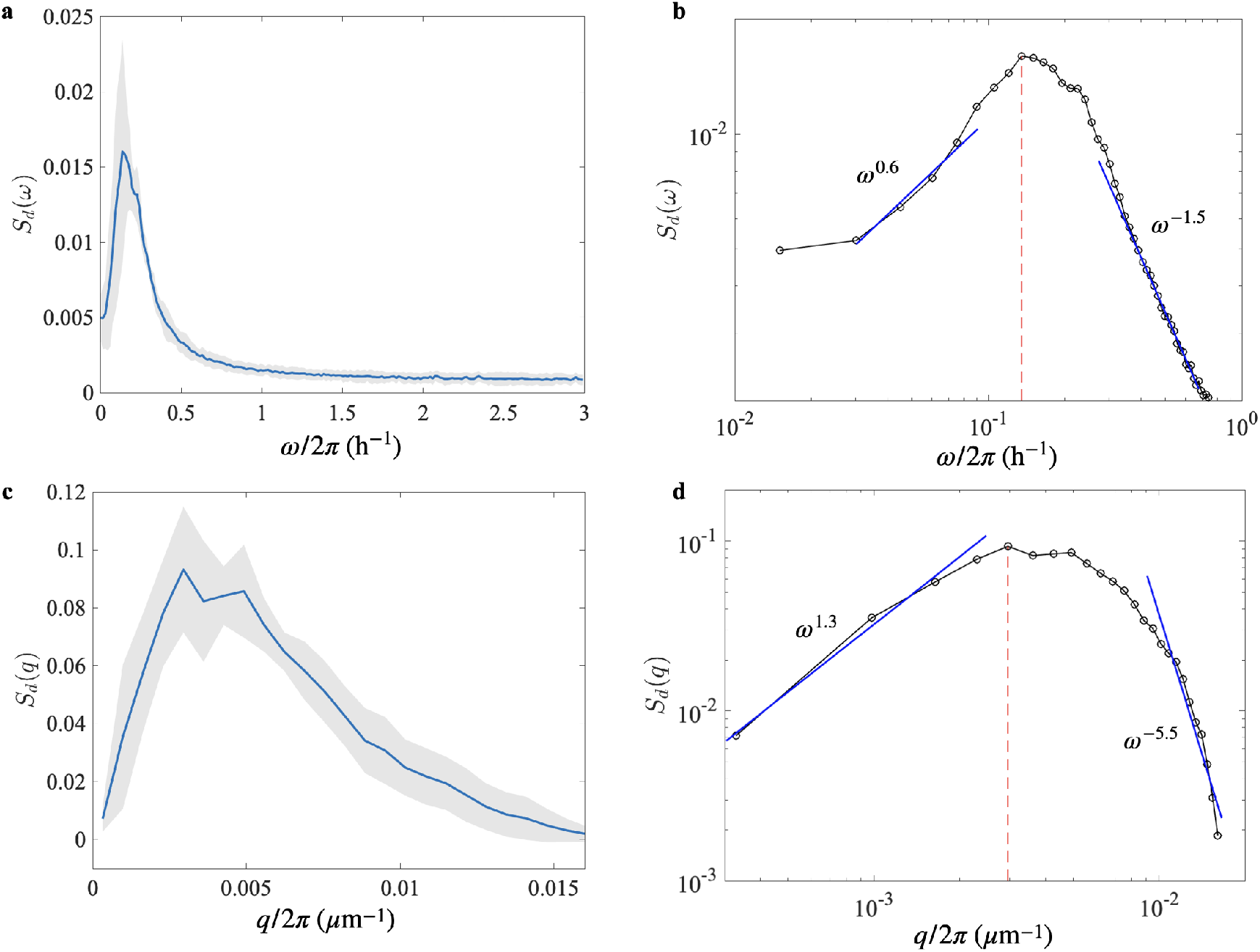
Power spectra (a) *S_d_*(*ω*), (c) *S_d_*(*q*) and the corresponding log-log plots (b) and (d) for pulsatory cellular movements in MDCK epithelial monolayers on non-patterned surfaces. The nature of the power spectra is similar to that obtained for patterned surfaces in figs. 3c and 3e. The blue lines and the grey shaded region in (a) and (c) correspond to the mean and standard deviation, respectively, of *S_d_*(*ω*) and *S_d_*(*q*) obtained over five separate experimental repeats. The power-law fits for low and high values of(b) *ω* and (d) *q* are represented with the blue lines and the corresponding exponents are indicated.

## Appendix C: Reasoning behind the term involving *K_h_* in tissue stress

We provide an intuitive reasoning for including the term 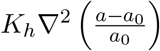 in tissue stress *σ_ij_* in Eq. 3. We first note that a cell has at least two modes of deformation. First is the in-plane area change *δa* = *a* – *a*_0_ without any gradients in tissue height *h.* For an incompressible tissue, *ah* = *a*_0_*h*_0_ due to which *δa*/*a*_0_ = −*δh*/*h*_0_. For the area deformation mode, the energy density of the tissue is *k*_1_(*a* – *a*_0_)^2^. The other mode of deformation in which there is no change in cell area, would necessary involve cell height gradients, and the energy density, to the lowest order, can be be written as *k*_2_|∇*h*|^2^. Thus the total energy of the tissue due to these two modes of deformation is

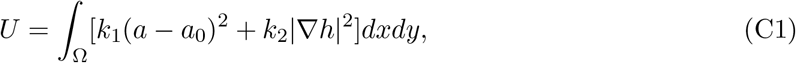

over the total region Ω of the tissue. We note that the apical and basal layers of the epithelium can have distinct mechanical properties [46]. Since spatial variation in tissue height *h* can lead to different deformations of the apical and basal layers, the difference in mechanical properties can further contribute to the tissue energy via the |∇*h*|^2^ term in Eq. C1. For small deformation of the tissue, the isotropic stress *σ* in the tissue will be obtained by taking the functional derivative of the tissue *σ* = *δU*/*δa*. After noting that ∇*h* = −*h*_0_/*a*_0_∇*a*, and renormalizing the values of the material coefficients *k*_1_ and *k*_2_ we get

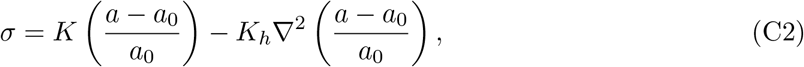

as written in Eq. 3. We note that in the absence of volume incompressibility of the monolayer, we cannot rigorously replace the ∇*h* term with ∇*a* as described above. Although, in such a case, we do not expect major qualitative changes in our model except for re-scaling of *K_h_,* the use of *∇a* term is, strictly speaking, a model approximation.

## Appendix D: Estimate for pattern length scale *l_0_*

We estimate the typical length *l*_0_ given in Eq. 10 by proposing the following experimental values for the parameters: substrate friction *η* ≈ 30 N · s · m^-1^ between the cells and the surface [47, 48], cell stiffness *Y* ≈ 100 pN · *μ*m^-2^ in terms of the typical force per unit area for cell-cell contacts [49], myosin homeostasis time *τ* ≈ 60 min [23], and myosin fluctuation correlation time along cell-cell junctions *τ_c_* ≈ 2 min [35]. Taking for cell area *a*_0_ ≈ 100 *μ*m^2^ and for cell height *h*_0_ ≈ 10 *μ*m, we can express *α* ≈ η × a_0_ and *K_h_* ≈ *Y* × *a_o_* × *h*_0_. Using these numbers we obtain an approximate length scale associated with the pulsating pattern *l*_0_ ≈ 100 *μ*m, which is consistent with the typical length scales measured experimentally.

