## Supplementary Information : Movie 1 for "Interplay between cell height variations and planar pulsations in epithelial monolayers"

**Movie 1:** Height fluctuations during monolayer pulsations. In the top, domain of MDCK-E-Cadherin-GFP cells on FN grid is shown at two z-planes: 2.5 $\mu$m and 8 $\mu$m. Scale bar, 100 $\mu$m. In the bottom, 3D surface view of cells where height is encoded as intensity is shown. Color bar shows the height values. Time in hh:mm.
